# REPP: A robust cross-platform solution for online sensorimotor synchronization experiments

**DOI:** 10.1101/2021.01.15.426897

**Authors:** Manuel Anglada-Tort, Peter M. C. Harrison, Nori Jacoby

## Abstract

Sensorimotor synchronization (SMS), the rhythmic coordination of perception and action, is a fundamental human skill that supports many behaviors, from daily repetitive routines to the most complex behavioural coordination, including music and dance (Repp 2005; Repp & Su, 2013). Research on SMS has been mostly conducted in the laboratory using finger tapping paradigms, where participants typically tap with their index finger to a rhythmic sequence of auditory stimuli. However, these experiments require equipment with high temporal fidelity to capture the asynchronies between the time of the tap and the corresponding cue event. Thus, SMS is particularly challenging to study with online research, where variability in participants’ hardware and software can introduce uncontrolled latency and jitter into recordings. Here we present REPP (Rhythm ExPeriment Platform), a novel technology for measuring SMS in online experiments that can work efficiently using the built-in microphone and speakers of standard laptop computers. The audio stimulus (e.g., a metronome or a music excerpt) is played through the speakers and the resulting signal is recorded along with participants’ responses in a single channel. The resulting recording is then analyzed using signal processing techniques to extract and align timing cues with high temporal accuracy. In this paper, we validate REPP through a series of calibration and behavioural experiments. We demonstrate that our technology achieves high temporal accuracy (latency and jitter within 2 ms on average), high test-retest reliability both in the laboratory (*r* = .87) and online (*r* = .80), and high concurrent validity (*r* = .94). We also show that REPP is fully automated and customizable, enabling researchers to monitor experiments in real time and to implement a wide variety of SMS paradigms. We discuss methods for ensuring high recruiting efficiency and data quality, including pre-screening tests and automatic procedures for quality monitoring. REPP can therefore open new avenues for research on SMS that would be nearly impossible in the laboratory, reducing experimental costs while massively increasing the reach, scalability and speed of data collection.

## Introduction

Sensorimotor synchronization (SMS) is a fundamental human skill that involves the temporal coordination of rhythmic movement with a predictable external event (Repp, 2005; Repp & Su, 2013). SMS requires individuals to precisely integrate visual or auditory perception with motor production, supporting a wide range of human behaviors. For example, the ability to entrain to an external auditory cue plays a key role in musical experiences across human cultures (Savage et al., 2015; Jacoby et al., in prep.) and has been linked to specific genotypes, suggesting an innate human sensitivity to rhythm (Niarchou et al., 2021). SMS has also been associated with the development of literacy skills, such as reading and speech (Carr et al., 2014; Flaugnacco et al, 2014; Ladányi et al., 2020; Tierney & Kraus, 2013), and various neurodevelopmental disorders, including attention deficit hyperactivity disorder (Noreika et al., 2013) and Parkinson’s disease (Bieńkiewicz & Craig, 2015).

Quantitative research on SMS dates back at least to 1886 (Stevens, 1886), but its popularity has increased considerably in recent decades (see Repp, 2005; Repp & Su, 2013, for reviews). SMS experiments can differ substantially in their implementation, using different production modes (e.g., finger tapping, clapping, or speaking), different stimulus domains (e.g., visual or auditory), and different experimental designs, including rate limits (London, 2002), perturbation studies (Repp 2002a), simulated partners (Repp & Keller, 2008), and transmission chains (Jacoby & McDermott, 2017; Ravignani et al., 2016). However, at their core, most SMS experiments consist of a relatively simple procedure: participants tap with their index finger to a rhythmic sequence of auditory stimuli. This procedure presents a methodological challenge: how to measure the asynchrony (or synchronization error) between the time of the tap and the corresponding cue event with high millisecond-level precision. To meet this challenge, previous studies have used various laboratory-based methods that rely on specialised software and hardware. For example, some studies have used external hardware to record responses and control auditory feedback (e.g., a MIDI percussion pad or keyboard connected to computer software), for example *FTAP* (Finney, 2001) and *Max-MSP* (Patel et al., 2005; Repp & London, 2005). Researchers have also proposed solutions that use the low-level timing hardware of Arduino microcontrollers, including *TapArduino* (Schultz & Vugt, 2016) and *TeensyTap* (Vugt, 2020). Another popular solution is *MatTAP*, a MATLAB based toolbox for dedicated data acquisition hardware (Elliot et al., 2009). Others have developed an iOS application for tapping experiments that takes advantage of specific hardware in mobile Apple devices (*Tap-It*, Kim et al., 2012). In previous work, we have used a simple, low cost, in-lab method that achieves high temporal fidelity by simultaneously recording the audio stimulus and tapping responses using a standard sound card with an audio loopback cable (Elliott et al., 2018; Jacoby & McDermott, 2017; see Experiment 2 for a description of this method).

Nevertheless, none of these methods are viable for performing SMS experiments in online settings, where researchers have very limited control over participants’ hardware and software. This lack of experimental control combined with the technical demands of SMS tasks makes studying SMS with online research a unique challenge. In particular, SMS experiments performed online, such as tapping on the spacebar or mouse in synchronization to an external beat, can introduce all kinds of delay in latency and jitter into the recorded timestamps (Anwyl-Irvine et al., 2020; Bridges et al., 2020). Latency refers both to the time gap between a participant pressing a key and the device registering the keypress, and the time interval between initiation of audio playback and the physical start of the sound. It is often related to issues concerning scan rates, device drivers, internet connections, operating system variability, and sound card start-up latencies. Jitter is closely related and refers to the variation in latency. It can be either introduced in each tapping onset or across tapping trials (e.g., operating systems usually process each keyboard stroke with different temporal latencies). These inaccuracies can be in the order of 60 to 100 ms and can vary considerably between platforms, browsers, and devices (Anwyl-Irvine et al., 2020). Thus, measuring participants’ asynchronies in online settings with high precision is currently unfeasible.

Another important source of noise in online experiments is altered participant behavior compared to laboratory settings (e.g., Clifford & Jerit, 2014). This can be challenging in SMS tasks because they usually require participants to pay close attention to the task and take a large number of trials per session. There is also a higher risk of fraudulent responders (Ahler et al., 2019; Crump et al., 2013), including both computer ‘bots’ and non-serious respondents, such as participants who do not tap at all or tap at a regular rate irrespective of the external auditory cue. When performing online research on SMS, it is therefore important to rapidly analyse experimental trials and monitor performance in real time, providing feedback to participants and excluding fraudulent responders.

Here we present REPP (Rhythm ExPeriment Platform), a novel technology for measuring SMS in online experiments that can work efficiently using hardware and software available to most participants online, specifically standard laptops with working speakers and microphones. To address core issues related to latency and jitter, REPP uses a free-field recording approach: the audio stimulus is played through the laptop speakers and the original signal is simultaneously recorded with participants’ tapping responses using the built-in microphone (Figure 1A). The success of this method relies on a simple observation: although the initial onset in a recording is hard to control due to the interplay between the sound card and operating system, once a sound card starts recording, it registers all subsequent sound events as audio samples encoded with high precision with respect to the beginning of the recording. Thus, by using a single audio recording to simultaneously capture the stimulus and tapping onsets, we can remove the most significant sources of delay in both response and presentation latencies. We then apply audio filtering and other signal processing techniques to the resulting audio recording to split the different components of the recording into separate channels and therefore extract the stimulus and tapping onsets with reliable timing. Finally, we use custom markers with known temporal locations to unambiguously identify the position of the tapping and stimulus onsets in the audio recording, allowing a precise alignment to measure participants’ asynchronies. REPP can be executed rapidly in real time and is fully customizable, enabling researchers to adapt the code to support a wide range of SMS paradigms in online and laboratory settings. In this paper, we aim to validate REPP in a series of experiments while demonstrating how to best implement it in online studies to ensure high data quality.

**Figure 1.**
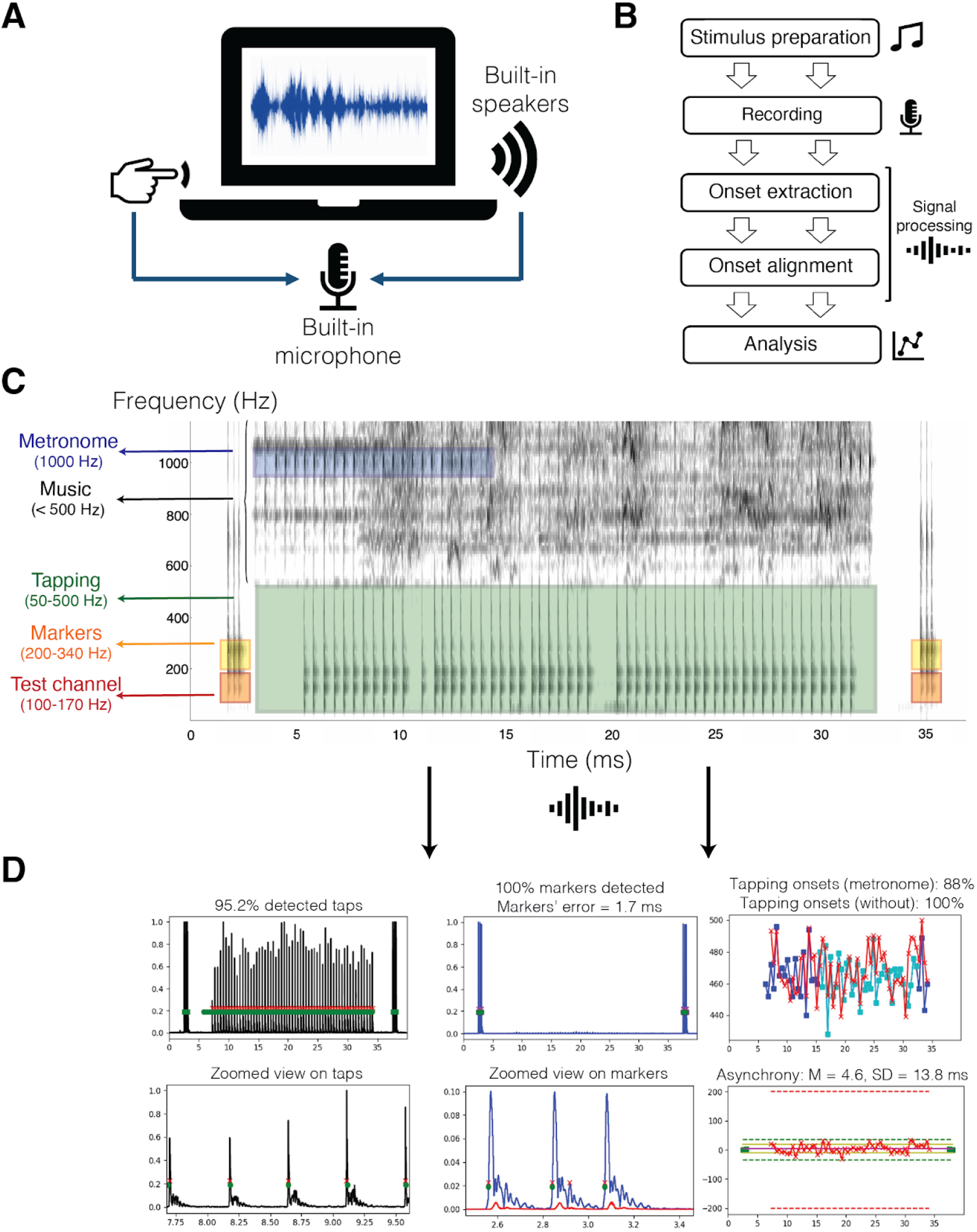
REPP: A robust cross-platform solution for online SMS experiments. (A) REPP uses a free-field recording approach that can work efficiently using standard hardware and software available to most online participants. (B) REPP comprises five main steps. (C) Example of a recording using REPP in a trial of beat synchronization to music. REPP uses a unique frequency range for each audio element in the recording: metronome (blue), tapping (green), markers (yellow), and test channel (red). (C) Output of the performance analysis after the signal processing steps, including the number of detected tapping onsets, detected markers, and mean and *SD* of asynchrony.

This paper continues with an overview of REPP. We then present a series of calibration and behavioral experiments demonstrating key aspects of this technology: temporal accuracy (Experiment 1), test-retest reliability and concurrent validity in the laboratory (Experiment 2), test-retest reliability in a larger scale online experiment, as well as methods for ensuring high data quality while minimizing costs (Experiment 3), and the ability to replicate a more complex tapping paradigm online (Experiment 4). Finally, we discuss the limitations of REPP and implications for future work on SMS. We will release REPP as a free and open-source framework with the published version of the paper.

### Overview - REPP

REPP can be organised around five main steps: (i) *stimulus preparation*, (ii) *recording phase*, (iii) *onset extraction*, (iv) *onset alignment*, and (v) *performance analysis* (see Figure 1).

REPP takes two inputs: an audio file with the stimulus (e.g., a metronome or a music clip) and a list of the corresponding stimulus onsets. Prior to performing an experiment, the stimulus must be prepared to be used efficiently in the subsequent steps. In particular, we first filter the stimulus to avoid any overlap with the frequency range that will be occupied by participants’ tapping responses (e.g., 50-500 Hz). To unambiguously identify the position of the tapping and stimulus onsets in the audio recording, we then add custom markers with known temporal locations to the beginning and end of the stimulus (see Figure 1C). These markers are designed to be robustly detected across participants’ hardware and software, including cases of noise-cancellation technologies and noisy recordings. In particular, we play the markers at low frequencies (typically 200-340 Hz) and nearly maximum volume.

In the recording phase, REPP uses a free-field recording approach: we play the prepared stimulus through the laptop speakers and simultaneously record the resulting audio signal along with the participant’s tapping response using the built-in microphone (see Figure 1A). This method returns an audio file where both the audio stimulus and tapping onsets are mixed in the same channel. The next step therefore applies signal processing techniques to split the mono recording into separate channels. REPP uses a unique frequency range for each relevant audio element in the recording, including a marker range and a tapping range (see Figure 1C). The tapping range is determined by the acoustic spectrum of the sound produced by participants’ mode of tapping. In our procedure, participants tap with their index finger on the surface of their laptop, producing a crisp sound with a significant part of its energy between 80 and 500 Hz. Since we have previously filtered the audio stimulus to avoid any overlap with this tapping range, we can efficiently extract the tapping signal from the raw recording by using bandpass filters with cut-off frequencies set to these ranges. Similarly, we use a specific range to filter and identify the marker locations (i.e., the same range used to generate the marker sounds, e.g., 200-340 Hz). In addition, to enhance the markers’ extraction procedure, we use a signal cleaning heuristic that baselines the amplitude of the filtered markers channel against a test channel set to a lower frequency range (Figure 1C). The rationale here is that tapping sounds will have similar energy within the markers and test channels, whereas marker sounds will have all energy in the markers channel and nearly no energy in the test channel. Thus, we can be sure that the detected signal corresponds to the markers and not to other sources of noise (e.g., tapping signal or background noise). This method also allows us to increase the signal-to-noise-ratio only in those areas containing the markers. Finally, REPP applies a simple onset extraction algorithm to the filtered tapping and markers channels to detect all samples exceeding a relative threshold (Elliott et al., 2018), returning a vector of extracted tapping onsets and extracted marker onsets.

The next challenge consists of aligning the extracted taps to their position in the audio stimulus. Since the sound card guarantees that all events are recorded with high precision with respect to the beginning of the recording, we can use the first detected marker as a single frame of reference to align the stimulus and tapping response. The other markers can be used to assess REPP’s timing performance in each trial and exclude fraudulent respondents or participants with incompatible hardware or software (see *Failing Criteria* in Appendix A). Importantly, by relying on the markers’ accurate timing, we do not need to extract the stimulus onsets from the recorded signal, which can be challenging in online studies due to noise-cancellation technologies and interference from other audio elements (e.g., participants’ tapping response and background noise). Instead, we use the list of stimulus onsets provided in the stimulus preparation step, allowing us to remove the audio stimulus from the actual recording and therefore minimize any interference with other elements in the signal processing pipeline. This method can also support SMS experiments using “virtual” onsets that are not clearly defined in the audio signal, such as when working with music (Colley et al., 2018; Dannenberg & Wasserman, 2009; Patel et al., 2005; Repp, 2002b). The output of this step is a list of re-aligned stimulus onsets and re-aligned tapping onsets. In the last step, we calculate several metrics to assess the performance of REPP and measure participants’ tapping accuracy, such as mean and standard deviation of the asynchrony (see Figure 1D).

### Validation Experiments

A total of four experiments were conducted to validate REPP and show how it can be implemented in online experiments to produce high-quality data. In Experiment 1, we assess the timing accuracy of REPP using an independent calibration system. In Experiment 2, we assess REPP’s test-retest reliability and concurrent validity when measuring individual differences in SMS in the laboratory. In Experiment 3, we assess the test-retest reliability of REPP with a larger sample of participants recruited online and also provide suggestions to reach high data quality while minimizing recruiting costs. Finally, Experiment 4 assesses REPP’s ability to implement more complex SMS paradigms in online settings, namely, transmission chain experiments designed to measure perceptual priors on rhythm. Additional methods and demographic information are presented in Appendix A.

To measure tapping accuracy, we followed common practices established in previous tapping studies (Repp, 2005). Asynchronies were defined as A_n_ = R_n_ – S_n_, where R_n_ denotes a response onset and S_n_ denotes a stimulus onset in a given tapping trial. We then computed the mean asynchrony and standard deviation (*SD*) of asynchrony. Throughout the experiments, we report the *SD* of the asynchrony, as it provides a more consistent measurement of tapping accuracy than mean asynchrony. Typically, the mean asynchrony is negative due to a human tendency to anticipate taps by a few tens of milliseconds when synchronizing to an external cue event (Repp, 2005). The mean asynchrony is also more influenced by tapping task, production modality, and auditory feedback biases compared with the *SD* of the asynchrony. However, we obtain very similar results when repeating the main analyses using mean asynchrony (see Appendix B).

### Experiment 1 - Timing Accuracy

This experiment assessed the timing accuracy of REPP by comparing its performance with a ground-truth recording obtained from an independent calibration system. The experiment was divided in three parts. Part 1 and 2 were large-scale validation experiments aimed to extensively test the timing accuracy of the audio stimulus and tapping response, respectively. Part 3 was smaller in terms of the number of data points but aimed to test all components of REPP together (i.e., markers, stimulus, and tapping response).

REPP’s timing performance was tested in the laboratory against an independent calibration system. This allowed us to measure the cue events (either tapping or stimulus onsets) separately in the two systems, providing an upper bound on inaccuracies of both REPP and the independent calibration system. Based on an established method previously used in our work (Jacoby & McDermott, 2017), we used a calibration system that offers a simple solution for measuring the ground-truth recording of REPP (see Figure 2A). We tested two variants of this system: one where participants tap on a tapping sensor (part 2) and another where participants tap on the surface of the laptop (part 3). In general, the independent calibration system uses two external synchronized devices to record the stimulus and tapping signals as soon as they are produced by the laptop speakers and finger tap, respectively. Since both the stimulus and tapping signals produce a highly precise sound wave, we can then apply a simple onset extraction algorithm to precisely identify the location of the onsets at the earliest possible moment. To record the audio stimulus (both the markers and stimulus onsets), we directed a Shure SM58 microphone to the laptop speakers being tested. To record the tapping onsets, we used a custom made tapping sensor device. The sensor consisted of a soft pad with earbuds installed inside (Apple EarPods). The earbuds offer a low-sensitivity microphone that is well-suited to precisely detect touch on the surface of the sensor while being insensitive to external noises and minimizing auditory feedback. The tapping sensor was placed next to the laptops’ built-in microphone to capture the sound of the finger tapping. Both the microphone and tapping sensor were connected to a Focusrite Scarlett 2i2 USB sound card to record the signal on a separate MacBook computer running *Ableton Live 10* Software, saving the resulting recording as a wave file.

**Figure 2.**
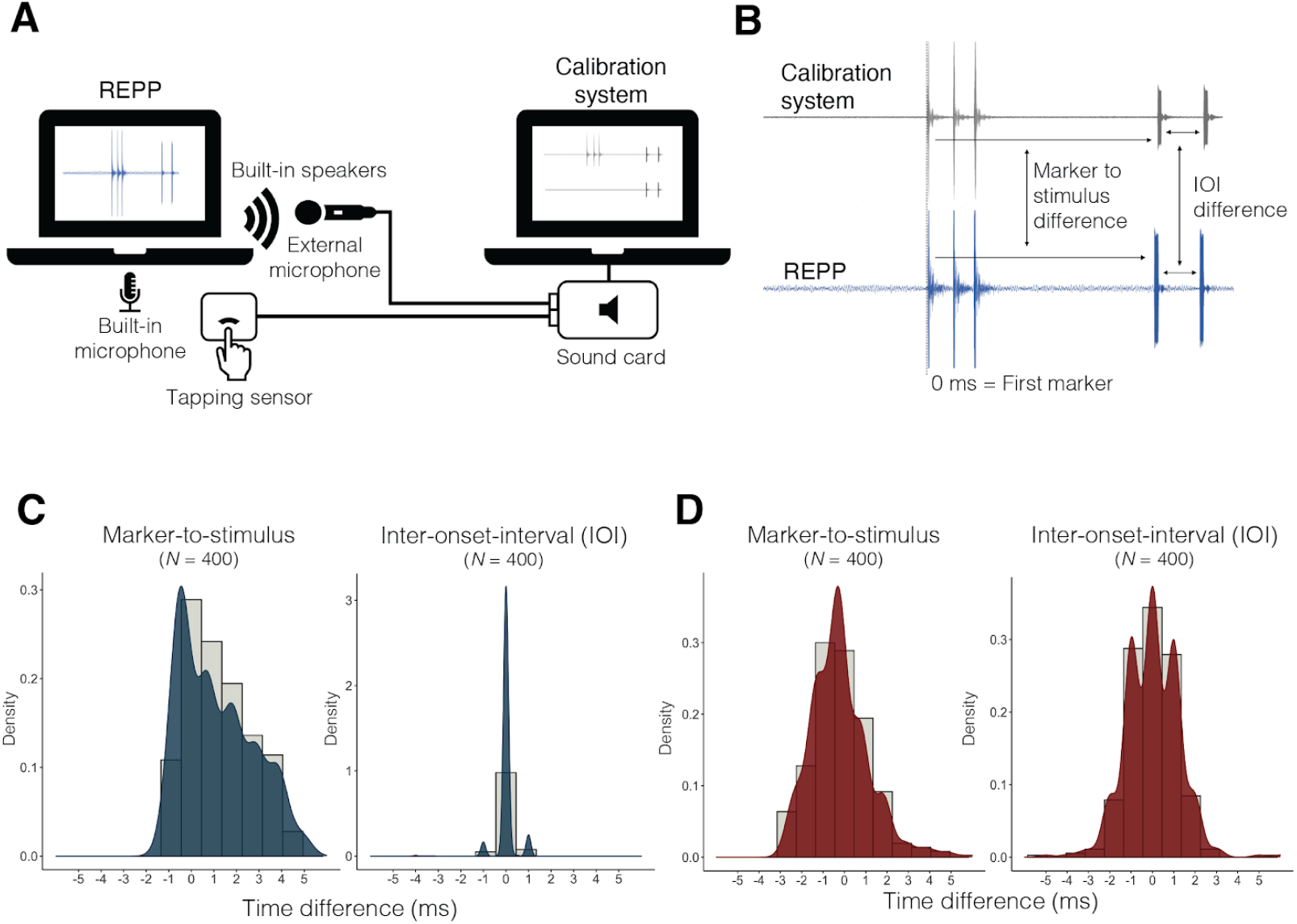
Results of Experiment 1 : Timing accuracy. (A) External calibration system used to measure REPP’s timing accuracy. (B) Example of the beginning of a 500 ms IOI trial recorded in the two systems, showing the three marker sounds placed at the beginning of the stimulus and the two first metronome clicks. The figure illustrates the two alternative measures to assess timing accuracy: the difference between the first marker and the stimulus (marker-to-stimulus), and the inter-onset interval. Note that the calibration system has two input channels (external microphone and tapping sensor) but we combine them in the figure for simplicity. (C) Distribution of the difference between the time the stimulus onsets (metronome clicks) were produced and the time they were detected by REPP. *N* refers to the total number of tested onsets. (D) Distribution of the difference between the time the physical taps were produced and the time they were detected by REPP. *N* refers to the total number of tested onsets.

To validate the timing accuracy of the audio stimulus (part 1), REPP was programmed to produce 100 isochronous metronome clicks at four different inter-onset intervals (IOIs): 250 ms, 500 ms, 750 ms, and 1000 ms. No finger taps were produced for this part and only the audio stimulus was recorded using the external microphone (Figure 2A). To validate the timing accuracy of the tapping response (part 2), the same trials were produced and a researcher tapped in time to the clicks, resulting in four trials of 100 taps at four different IOIs: 250 ms, 500 ms, 750 ms, and 1000 ms. The tapping response was recorded using the tapping sensor and the audio stimulus was recorded using the external microphone (Figure 2A). In the third part, to validate the timing accuracy of all components of the system together (i.e., markers, stimulus, and tapping response), REPP was programmed to produce 20 isochronous metronome clicks at two IOIs: 500 ms and 1000 ms. This time, the researcher tapped on the surface of the laptop in anti-phase, so the stimulus and tapping onsets could be unambiguously distinguished in the recording. Both stimulus and tapping onsets were recorded with the same external microphone. The recording was then separated into three channels (i.e., markers, stimulus, and tapping response) and manually cleaned to only contain the corresponding elements in each channel (e.g., stimulus onsets in the stimulus channel, tapping onsets in the tapping channel).

The results of the timing accuracy analysis in all validation parts are reported in Table 1. The average latency and jitter of REPP was within 2 ms and similarly accurate for all components of the system: markers, audio stimulus, and tapping response. Timing accuracy was computed as the difference between the time the stimulus or tapping onsets were produced (measured using the external calibration system) and the time they were detected by REPP, using the first detected marker as the single frame of reference (Figure 2B). We used two alternative measurements to calculate timing accuracy: marker-to-stimulus or marker-to-tap (i.e., the interval between the first marker onset and each subsequent onset in the audio or tapping signal), and inter-onset-interval (i.e., the interval between onsets). Figure 2C and 2D show the distribution of the time difference between the stimulus onsets (part 1) and tapping onsets (part 2) measured in the two systems, confirming that REPP’s timing accuracy is high and consistent.

**Table 1.** Timing accuracy results. Part refers to each validation experiment: part 1 (only stimulus), part 2 (only tapping response), and part 3 (stimulus and tapping response together). *N* refers to the total number of tested onsets.

| Part |  | *N* | Min-max | Latency (*SD*) | min-max | Latency (*SD*) |
| --- | --- | --- | --- | --- | --- | --- |
|  |  |  | *Relative to first marker* |  | *Inter-onset-interval* |  |
| 1 + 2 | Markers | 40 | -.08 - 4.9 | 1.85 (1.76) | - | - |
| 1 | Stimuli | 400 | -1.3 - 4.9 | 1.15 (1.61) | -4 - 1 | .02 (.39) |
| 2 | Tapping | 400 | -2.8 - 5.9 | -.2 (1.37) | -5 - 8 | .03 (.13) |
| 3 | Markers | 10 | 0 - 1 | .58 (.48) | - | - |
| 3 | Stimuli | 40 | -.3 - .9 | -.03 (.23) | 0 - 1 | .04 (.17) |
| 3 | Tapping | 40 | -3.1 - 1 | -1.04 (.89) | -4 - 2 | -.02 (1.13) |

### Experiment 2 - Reliability and Concurrent Validity

Experiment 1 showed that REPP can measure tapping and stimulus onsets with high temporal accuracy. In Experiment 2, we aimed to examine whether REPP can reliably measure derived psychological quantities such as a particular individual’s tapping accuracy. Specifically, we assessed the test-retest reliability of REPP and compared its performance against a completely independent method: a well-established method previously used in the laboratory to measure SMS with high precision (Elliot et al., 2018).

To assess test-retest reliability, the same group of participants (*N* = 20) performed a short battery of tapping tasks two times in each method, using the following sequence: method 1 (pre), method 2 (pre), method 1 (post), method 2 (post). Half of the participants started the experiment using REPP, whereas the other half started using the independent in-lab method. To assess concurrent validity, we correlated the participants’ overall tapping performance in the two methods. Participants completed the experiment in a quiet testing room with two tables, one for each method.

The independent in-lab method consisted of a loop-back setup to measure participants’ asynchronies with high temporal fidelity (see Figure 3A). This method has been extensively used in the laboratory and field research on SMS (Elliot et al., 2018; Jacoby & McDermott, 2017; Jacoby et al., in prep). The loop-back setup consists of a cable connected to a sound card to simultaneously record the input signal from the headphones and the output signal from a tapping sensor with nearly zero latency. We used professional headphones (i.e., Sennheiser HD 280 Pro headphones) to deliver the stimulus, and the custom made tapping sensor described in Experiment 1 to record participants’ tapping response with high precision. Both the headphones and tapping sensor were connected to an external sound card (Focusrite Scarlett 2i2 USB) using the loop-back setup described above (Figure 3A). The sound card was connected to a MacBook via USB. The tapping tasks were implemented using MATLAB, mirroring the experimental procedure used in REPP. To detect the stimulus and tapping onsets at the earliest possible moment, we used the same onset extraction algorithm described in Experiment 1 (Jacoby & McDermott, 2017).

**Figure 3.**
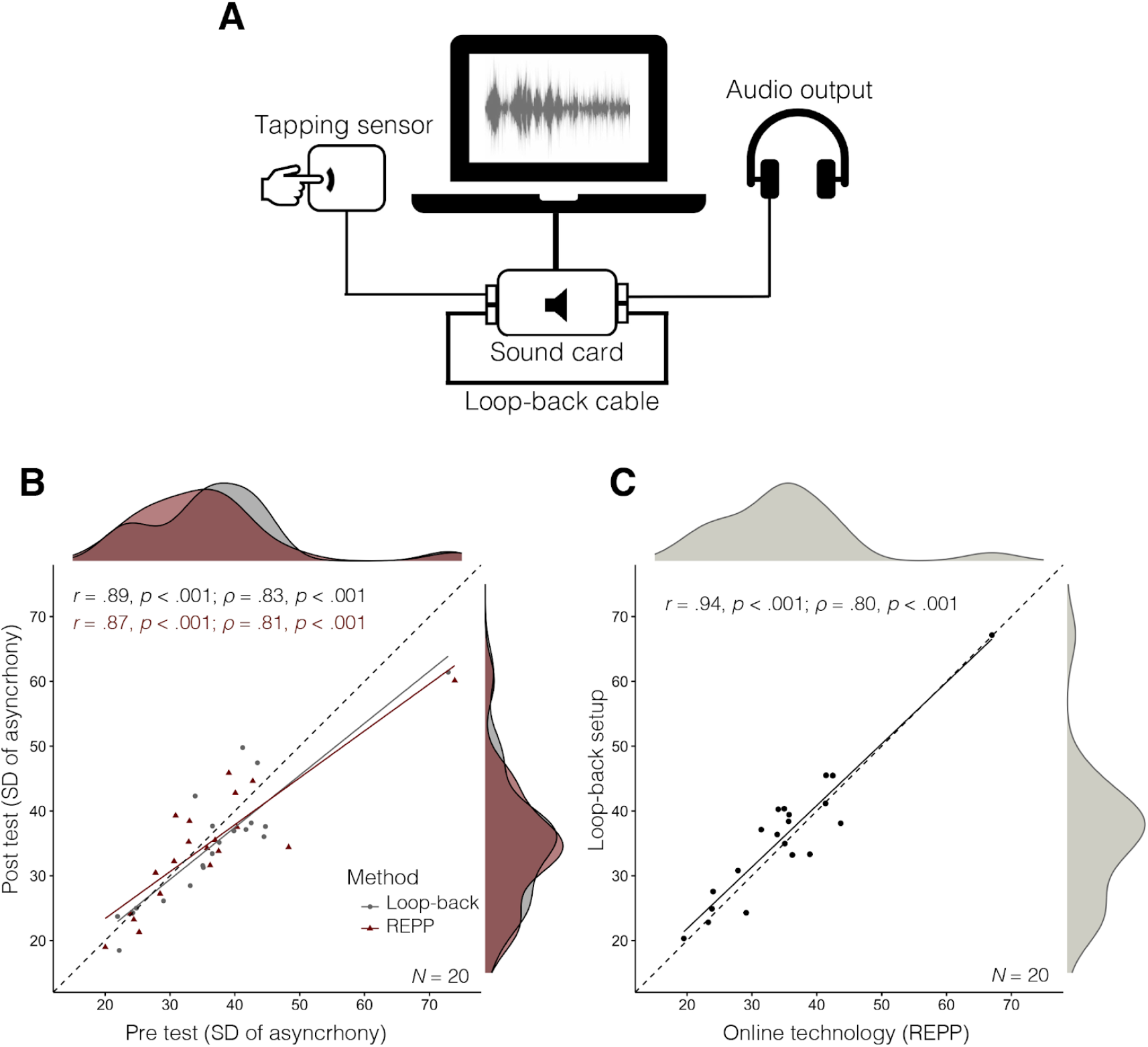
Results of Experiment 2 : reliability and concurrent validity. (A) Loop-back setup: independent in-lab method using a loop-back cable to measure participants’ tapping asynchronies with high temporal fidelity (Elliot et al., 2018). (B) Test-retest reliability in the two methods. (C) Concurrent validity: correlation between the overall tapping performance measured in the two methods.

The materials and experimental procedure were identical in the two methods (see *instructions* in Appendix A). Before starting the main tapping task, participants performed a practice phase to get familiar with each method (see *practice phase* in Appendix A). The main tapping tasks consisted of a short battery of tapping trials (8-10 minutes long approximately) using two common paradigms in the tapping literature (Repp, 2005; Repp & Su, 2013): isochronous tapping and beat synchronization to music. The isochronous tapping consisted of four 30-second long trials of isochronous tapping to a metronome sound (two with IOIs of 800 ms and two with the IOIs of 600 ms). The presentation order was fixed, using the following sequence: 800 ms, 600 ms, 800 ms, and 600 ms. The beat synchronization task consisted of four 30-second long excerpts of music from two distinct music genres with different style, tempo, and tapping difficulty, also with fixed order of presentation (see *Beat Synchronization task* in Appendix A).

The results of Experiment 2 are plotted in Figure 3. To examine test-retest reliability, an aggregated performance score was calculated for each participant in each test (pre and post) and method by averaging their tapping performance in the two tapping tasks (i.e., isochronous tapping and beat synchronization to music). The test-retest correlation in REPP was high (*r* = .87; *ϱ* = .81) and similar to the one achieved by the independent loop-back setup (*r* = .89; *ϱ* = .83; Figure 3A). We further examined test-retest reliability by calculating the intraclass correlation coefficient (ICC; Shrout & Fleiss, 1979). Following the recommendations of Koo and Li (2016), ICC estimates and their 95% confidence intervals were calculated based on single-rating, absolute-agreement, 2-way mixed-effects models (ICC3). We found a good ICC in both REPP (ICC = .86, 95% [.72, .93]) and the loop-back setup (ICC = .89, 95% [.77, .95]). This ICC is comparable to the values reported in previous work assessing the test-retest reliability of similar rhythmic production tasks (Bégel et al., 2018). Moreover, an ANOVA confirmed that participants’ mean tapping performances were similar across test-retest conditions and tapping tasks (all *p*-values > .05). Finally, we found that REPP has a high concurrent validity (*r* = .94 and *ϱ* = .89), as indicated by the correlation between the overall tapping performances (averaging over both test and retest) measured by the two methods (Figure 3B). In conclusion, the converging evidence of these analyses is that REPP produces reliable estimates to measure individual differences on SMS in a way that is consistent with the results produced by a completely independent method.

### Experiment 3 - Online Demonstration

Having demonstrated that REPP achieves high temporal accuracy (Experiment 1) and test-retest reliability in the laboratory (Experiment 2), this experiment aimed to show that the technology can work in practice in an online setup that is similar to a large-scale data collection process. We also provide suggestions to ensure high data quality while enabling realistic data collection, in particular concerning pre-screening tasks and feedback based on recording quality and tapping performance. Participants were recruited from Amazon Mechanical Turk (see *Participants* in Appendix A) and performed the same battery of tapping tasks used in Experiment 2. In a total of six experimental batches (8 to 10 hours each), we collected valid tapping data for 226 participants.

We used two pre-screening tasks to ensure high data quality while minimizing recruiting costs (see *Pre-screening Tests* in Appendix A). First, we used an attention test to determine whether participants were paying attention to the instructions. Participants who failed the attention test were excluded from the experiment. Second, we used a recording test to determine whether participants were using hardware and software that were not compatible with REPP, such as malfunctioning speakers or microphone, or the use of strong noise-cancellation technologies. Participants who did not pass the recording test were also excluded from the experiment. To assess the efficacy of the recording test in comparison to the attention test on its own, we only used the recording test in half of the participants.

Before the main tapping tasks, participants were instructed on several key aspects concerning the proper functioning of REPP, including instructions about the technical requirements and tapping procedure for the experiment, a volume calibration test, and a tapping calibration test (see *Instructions* in Appendix A). Next, participants undertook a practice phase consisting of four trials of isochronous tapping to a metronome sound (see *Practice Phase* in Appendix A). After completing the practice phase, the four audio recordings were analyzed in real time using a failing criteria designed to identify and fail trials where participants used incompatible hardware and software, or where participants did not tap as indicated in the instructions (see *Failing Criteria* in Appendix A). Those participants who failed two or more trials were excluded from the experiment. After the practice phase, participants started with the main experimental task, which consisted of the same battery of tapping tasks employed in Experiment 2 (i.e., four trials of isochronous tapping and four trials of beat synchronization to music, 30 seconds long each). Participants repeated the same battery of tapping tasks a second time in order to measure test-retest reliability.

The results of Experiment 3 are visible in Figure 4. For measuring test-retest reliability, we only consider participants who provided at least one valid tapping trial for each stimulus in each tapping task and test-retest condition (*N* = 166). Test-retest analyses were performed using the same procedure described in Experiment 2. Results indicated a high test-retest correlation when using REPP to measure participants’ tapping performance online (Figure 4A; *r* = .80 and *ϱ* = .81), also confirmed by an intraclass correlation analysis (ICC = .82, 95% [.77, .86]). Moreover, participants’ tapping performance was similar across test-retest conditions in the two tapping tasks, as indicated by two paired samples *t*-tests with test condition as the independent variable and tapping performance in each tapping task as dependent variables (all *p*-values > .05). As a measure of convergent validity, we further examined whether participants’ tapping performance was related with their self-reported levels of musical training. We calculated an aggregated tapping performance score (averaging over test and retest in the two tapping tasks) for all participants who provided good tapping (*N* = 226). Musical training was measured using a reduced version of the Gold-MSI musical training factor (Müllensiefen et al., 2014). Replicating a recurring finding in the literature (e.g., Niarchou et al., 2021; Repp, 2010; Thompson et al., 2015), we found a significant negative correlation (*r* = −.32, *p* < .001) between tapping variability and self-reported musical training (Figure 4B), indicating that more musically trained participants were better at synchronizing to an external beat. To explore the robustness of our technology across operating systems and laptop models, we compared the marker detection accuracy (i.e., the delay between the known marker locations and the detected marker onsets) across the two most common operating systems, Windows and macOS (Figure 4C). Overall, trials recorded in macOS computers achieved slightly better temporal accuracy (*M* = 1.48, *SD* = .68) than trials recorded in Windows (*M* = 1.74, *SD* = .85), *t*(2812) = 9.36, *p* < .001. A small difference in this direction is not surprising: macOS computers are typically better equipped for delivering sound and recording audio than Windows computers, which also tend to exhibit greater variability in hardware.

**Figure 4.**
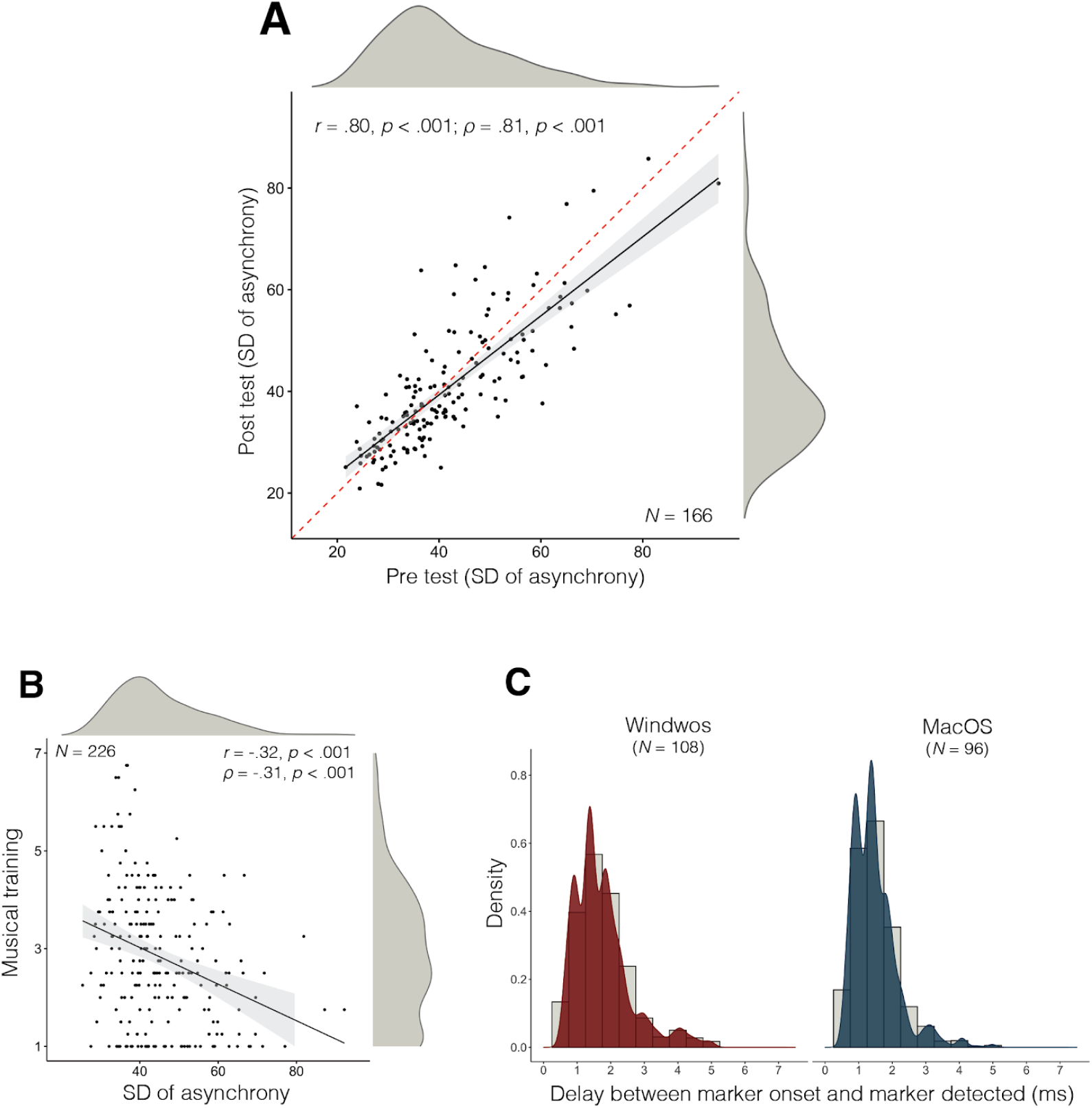
Results of Experiment 3: Online demonstration. (A) Test-retest reliability of REPP when measuring participants’ tapping performance online. (B) Convergent validity: correlation between overall tapping performance and participants’ musical training. (C) Estimated markers’ error in Windows and macOS computers. *N* indicates the number of participants using each Operating System, but we plot the data in all tapping trials (2,814 in total).

### Experiment 4 - Iterated Tapping (Online Replication)

This experiment tested REPP’s capacity to support a relatively complex tapping paradigm: estimating perceptual priors for simple rhythms via iterated reproduction of random temporal sequences, following a paradigm from Jacoby and McDermot (2017).

Participants (*N* = 157) were presented with random “seed” rhythms (clicks separated by random time intervals) and asked to reproduce the rhythms by tapping. The initial seed was randomly sampled uniformly from a two-dimensional triangular simplex comprising of three-interval rhythm that spans a constant duration (2,000 ms in this experiment), also known as “chronotopological map” (Desain & Honing, 2003). Every point on this triangle represents a unique three-interval rhythm with a fixed duration (and therefore two degrees of freedom). The stimulus presented to participants was generated by repeating the corresponding three-interval seed pattern 10 times. For each reproduction, we programmed REPP to extract the tapping onsets and average the inter-response interval across the 10 repetitions. The resulting averaged reproduction was then substituted for the seed and used to generate on the fly the stimulus for the next iteration. The process was iterated five times. In particular, each participant approximately completed four chains with five iterations derived from a single random seed (i.e., 20 trials), yielding to 507 within-participant chains^1^. Each iteration used the averaged reproduction from the previous trial. Over time, participants’ reproductions become dominated by internal biases and perceptual priors can be estimated by repeating this procedure multiple times (using multiple chains). We adapted the instructions, pre-screening tasks, and practice phase described in Experiment 3 (see Appendix A).

Figure 5 shows the results of the online replication using REPP. The original (laboratory) results are from the non-musicians Experiment 1 in Jacoby and McDermott (2017). To estimate the continuous distribution underlying the responses, we applied kernel density estimation to the last iteration’s data (Figure 5A). To clarify the structure of the final distribution, we superimpose symbols (crosses) at rhythms whose intervals are related by simple integer ratios, resulting in 22 simple ratios. It is visually apparent that the results of the original and replication maps are similar with only nuanced differences, showing that the online experiment can fully replicate an extremely complicated pattern of behaviour obtained in a controlled laboratory experiment. To quantify the similarity and differences between the original and replication experiments, we fitted a constrained 22-component Gaussian mixture model following the procedure used in Jacoby and McDermott (2017), where the resulting modes reflect the integer ratios common in Western music. We focus on the weights of the categories as they represent the category’s perceptual “strength” and were used in the previous study for group comparison. Jacoby and McDermott (2017) found that the correlation between groups of US participants who differed in musical experience was high (*r* = .79, *p* < .001), and significantly higher than the correlation between participant groups from different cultures (*r* = .19, *p* = .43, for US and Amazonian participants). In line with these findings, we found that the weights of the 22 components were highly correlated across the lab and online experiments (*r* = .77, *p* < .001).

**Figure 5.**
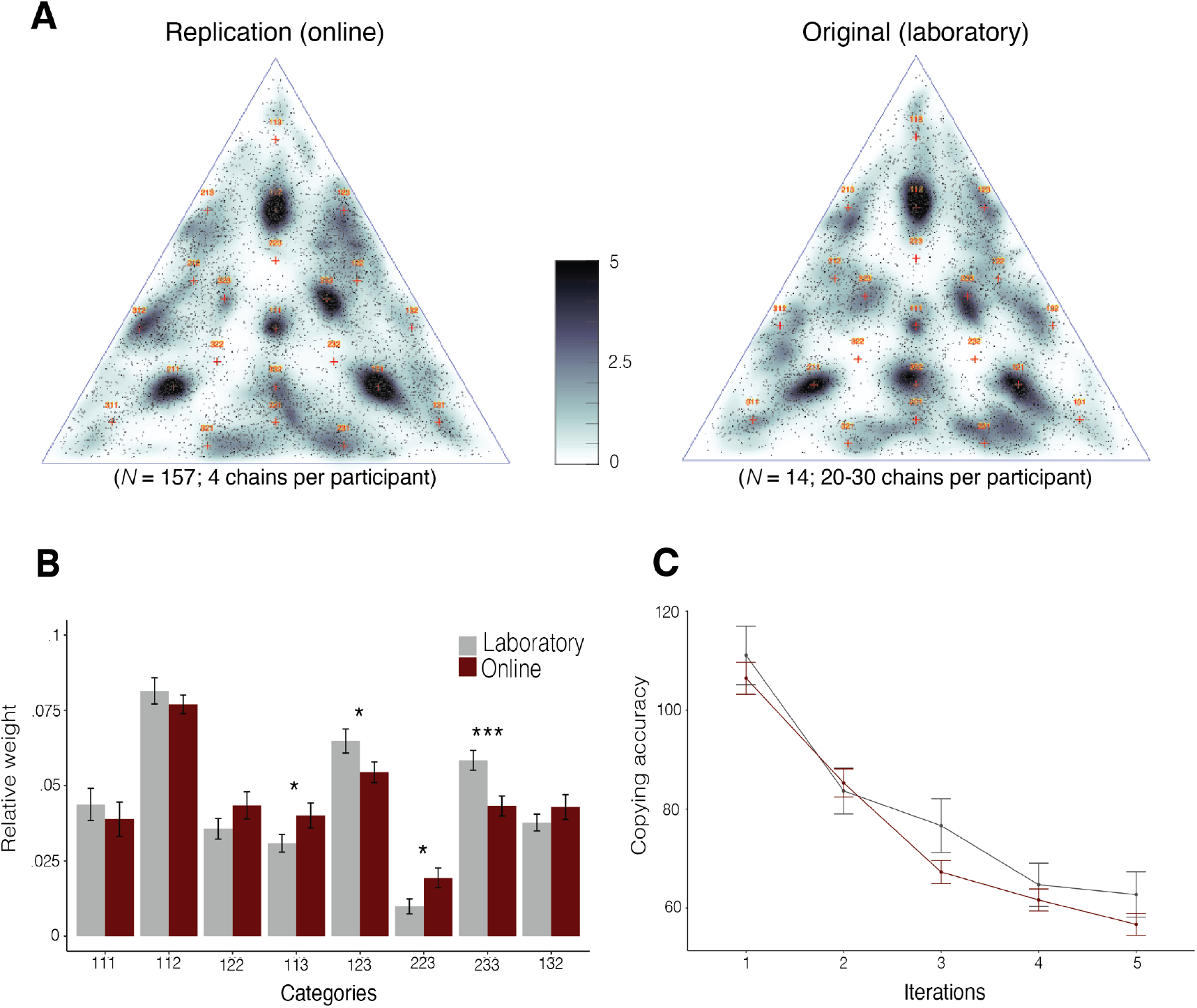
Results of Experiment 4: online replication of a transmission chain experiment. (A) Kernel density estimate of the continuous distribution underlying the data from iteration 5 in the original study and replication study online. Crosses plot simple integer ratio rhythms. (B) Weights of Gaussian mixture components assigned to eight main rhythm categories. (C) Copying accuracy (the distance between stimulus and reproduction) in the two experiments. Error bars represent confidence intervals on the weights (*SD* of the weight distribution derived from bootstrapping).

Figure 5B shows the fitted weights averaged over categories that are circular rotations of one another (e.g., 1:1:2 = 1:1:2, 1:2:1, 2:1:1), thus forming 8 categories. We average over the three rotations because of the apparent triangular symmetry for the results reported in the original study. We used bootstrapping (*N* = 1,000) to compute error bars and examine group differences on the eight categories. This comparison shows some small but significant differences in four categories (Figure 5B), perhaps due to group differences in either musical experience or cultural background (as participants in the laboratory and online may differ in these dimensions). Finally, we examined participants’ asynchronies and copying accuracy (the distance between stimulus and reproduction, which provides a measure of convergence speed and trial dynamics) in the two experiments. Copying accuracy over iterations was similar in the two experiments (Figure 5C), improving across iterations but not significantly different for the last two iterations (*t*(134) = 2.37, *p* = .07), suggesting that convergence is reached or nearly reached by the fifth iteration. Moreover, participants’ tapping variability (*SD* of asynchrony) across all tapping trials measured in the original laboratory experiment (*M* = 65.24, *SD* = 16.269) was nearly identical to the performance measured in the online replication experiment (*M* = 66.17, *SD* = 15.51), also confirmed by a Mann Whitney U Test (*U* = 1231, *p* = .46).

## Discussion

SMS is an active area of research with a long history of tapping experiments performed in the laboratory (Repp, 2005; Repp & Su, 2013). However, it currently lacks a robust method to precisely measure participants’ asynchronies in online experiments. In this paper, we presented REPP, a cross-platform solution for online SMS experiments that achieves high temporal accuracy and reliability while also being practical in terms of large-scale data collection. We plan to release this technology as a free and open-source framework alongside the journal version of the paper.

We validated REPP in a series of experiments. We first demonstrated that it achieves high temporal accuracy using an independent calibration system (Experiment 1). Based on the ground-truth recording, we estimated the average latency and jitter of REPP to be within 2 ms and similarly accurate for all elements in the system (i.e., markers, audio stimulus, and tapping response). In a laboratory experiment (Experiment 2), we then compared the test-retest reliability of REPP with a completely independent method that uses specialized equipment (i.e., a loop-back setup) to measure SMS with optimal temporal fidelity. We found that our technology achieves a high test-retest reliability (*r* = .87, *ϱ* = .81, ICC = .86) that is equivalent to the reliability obtained by the independent in-lab method (*r* = .89, *ϱ* = .83, ICC = .89). By correlating the overall tapping performance measured in the two methods, we also found that REPP has a high concurrent validity (*r* = .94 and *ϱ* = .80).

We then performed two experiments to show how REPP can work in practice in an online setup that is similar to a large scale data collection process. In Experiment 3, we confirmed that REPP has a high test-retest reliability using a larger sample of participants recruited online (*N* = 166; *r* = .80, *ϱ* = .81, ICC = .82). We also provided suggestions of tapping instructions and pre-screening tests to ensure high data quality in online experiments while minimizing recruitment costs (see Appendix A). In Experiment 4, we demonstrated that REPP is fully extensible and customisable, enabling relatively more complex tapping paradigms in online settings, such as transmission chain experiments where new stimuli are generated on the fly based on previous tapping responses. In particular, we replicated online (*N* = 157) a complex pattern of tapping behaviour previously obtained in a controlled laboratory experiment (Jacoby & McDermott (2017). The results confirmed that perceptual priors in US participants show peaks at rhythms with simple integer ratios. Together, these experiments demonstrate that our technology is well-equipped to support a wide variety of SMS experiments using standard hardware and software available to most online participants.

REPP has currently some limitations. First, it does not support real-time response feedback (Mates & Aschersleben, 2000; Finney & Warren, 2002). A possible solution would be to play the real time feedback with a Javascript audio process; this feedback may have compromised accuracy, but at least the feedback signal could be recorded and monitored with a variant of our technology. Developing this approach would however require significant additional work. Second, REPP relies on a stimulus preparation step that filters the audio stimulus to remove lower frequencies that would otherwise interfere with other aspects of the signal processing pipeline, such as the analysis of participants’ tapping response. This procedure decreases the perceived quality of complex auditory stimuli, such as music. However, we have shown that music stimuli can be perceived well to perform beat synchronization tasks after applying bandpass filters with cut-off frequencies below 800 Hz (Experiment 2 and 3).

Importantly, we learned that collecting good tapping data in online settings can necessitate a high exclusion rate, at least when using a large-scale recruiting strategy via Amazon Mechanical Turk. In Experiment 3, for example, a total of 727 participants began the online task. This includes anyone who accepted the experiment regardless of their intentions to take the task seriously or whether they met the technical requirements to provide good tapping data. Thus, we used a practice phase to familiarize participants with the task and exclude cases who could not provide good tapping data in the majority of trials. Note that we used relatively strict failing criteria to exclude trials based on whether the signal could be correctly recorded and whether participants produced a minimally acceptable number of tapping responses (see *Failing Criteria* in Appendix A). A total of 226 participants (31%) passed the practice phase and were able to provide good tapping data (a similar ratio was found in Experiment 4). The remaining 483 participants (69%) were excluded from the experiment and comprised a mix of fraudulent participants (e.g., computer bots or non-serious responders) and participants that did not meet the technical requirements of REPP, such as poor internet connection or incompatible hardware and software. For instance, common sources of failure were due to participants’ behaviour, such as not tapping at all, using desktop computers without built-in microphones, or performing the experiment with headphones instead of the laptop speakers. We also noticed that many participants did not follow the instructions to eliminate background noise, such as music or speaking, resulting in noisy recordings. An additional problem was the usage of remote desktops, which may be used by some participants to alter their geographical reported location. Since the remote desktop will open a microphone that is not physically connected to the computer of the participant, the technology is not able to record any signal. Furthermore, there were several cases where the technology failed due to laptops with low quality or malfunctioning speakers. The same occurred in laptops with strong noise-cancelling technologies, where the marker sounds are suppressed and cannot be detected in the signal. This last issue requires further investigation, as in theory it is possible to turn off noise cancellation manually in most devices, but the way to do so changes in different computer models and brands.

Naturally, the more demanding the online tasks, the higher the exclusion rate. In previous work, we found exclusion rates of only about 10% in Amazon Mechanical Turk experiments with minimal technical requirements, such as when using visual rating scales in the browser (Harrison et al. 2020). However, the exclusion rate increases when the experiment becomes technically more demanding. For example, a pre-screening test that requires participants to wear headphones to perform an auditory perception task produces an estimated exclusion rate of 36% (Wood et al., 2017), whereas performing online research with computer webcams can necessitate an exclusion rate of about 40% (Tran et al., 2017). In language production experiments that require participants to record themselves using a microphone to extract voice onset latencies, the exclusion rate can be around 60% (Vogt et al., 2021). Thus, the exclusion rate of REPP (~60-70%) is not unexpected when using a large-scale recruiting strategy via Amazon Mechanical Turk, as it is technically more demanding than previous paradigms. In particular, REPP can only work in SMS experiments when the marker sounds can be detected with high millisecond-level precision and participants take the task seriously (i.e., tapping with their index finger on the surface of their laptop in time to an auditory stimulus). A high exclusion rate is not particularly problematic when using online recruitment systems with large pools of active participants, but other modes of recruiting may require different strategies, such as when recruiting participants from special populations or using internal university systems. In these cases researchers can significantly reduce exclusion rates by using more relaxed failing criteria and taking more time to support participants and ensure they follow the instructions and meet the technical requirements (e.g., make sure they use the laptop built-in speakers with high volume, disable noise-cancelling technologies, and explicitly grant access to record in the browser). REPP can also be used in laboratory studies and field research with an exclusion rate of effectively 0%, as shown in Experiment 2.

Since exclusion rates may be high when using technical demanding tasks in online recruiting systems, such as Amazon Mechanical Turk or Prolific, we strongly recommend the use of pre-screening tests to determine whether participants will take the experiment seriously and meet the technical requirements to provide good tapping data. In Experiment 3, we analyzed the efficacy of two pre-screening tests, an attention test and a recording test (see Appendix A for a full description). We defined the exclusion rate of the pre-screening tests in terms of the proportion of participants who successfully passed the practice phase. Accordingly, we found that when using both pre-screening tests, 68% of the participants were able to pass the practice phase and deliver good tapping data in the experiment. In contrast, when only using an attention test without the recording test, the percentage was 31%. Thus, adding a recording test at the beginning of the experiment reduces the costs of recruiting participants online by nearly half. Such practices can also help maintain a good reputation in the online community; for example, both the attention test and recording test help eliminate a large proportion of the failure rate that comes from fraudulent participants, including computer bots and non-serious respondents (Crump et al., 2013). In addition to pre-screening tests, we encourage the use of data quality checks to monitor participants’ performance throughout the experiment. REPP computes several metrics to check the quality of a given recording, such as the number of detected markers or the time error between the known locations of the markers and the detected onsets, which provides a reliable measure of timing accuracy. We can also know how well participants are tapping by computing the ratio between the number of detected onsets and the number of stimulus onsets. Using these metrics, we provided feedback after participants’ completed the first tapping trial in the practice phase (Experiment 3 and 4), indicating whether their recording quality was sufficiently good and if it was not, suggesting ways to improve it for the subsequent trials. We encourage future research to explore these options further in order to increase recrurinement efficiency, such as providing more detailed feedback after tapping trials or nudging participants online to meet the technical requirements.

It is worth noting that REPP can be easily extended to support online experiments requiring precise timing of tapping response without any synchronization to an external stimulus. For example, unconstrained finger tapping paradigms where participants are not given an external stimulus but instead are asked to tap at their preferred rate (e.g., Collyer et al., 1994), or imitation experiments in which participants replicate a rhythm from memory (e.g., Ravignani et al., 2016). In these cases, researchers can skip the stimulus preparation and onset alignment steps, and simply use the parts of the pipeline that are directly related to the onset extraction procedure. We have successfully explored these options using a simplified method for several experiments that do not require stimulus-response synchronization, and support this variant in the code package associated with the journal version of this paper. Another simple extension of REPP is from finger tapping to other modes of production, including clapping, tapping on a table, or speech. We noticed that our technology works well also for clapping or tapping on a table, but adapting it to spoken utterances may require a modification to the parameters of the signal processing pipeline (see Experiment 6 in Jacoby & McDermott, 2017). Potentially, our technology could also be used to support online experiments requiring precise timing in domains other than rhythm perception and production. This includes any experiment measuring reaction times to auditory stimuli, such as auditory lexical decision tasks (e.g., Blumstein et al., 1982; Goldinger, 1996), priming paradigms using spoken words (e.g., Radeau et al., 1998), sounds (e.g., Schön et al., 1998), or music (e.g., Bharucha & Stoeckig, 1986, 1987), and experiments on temporal processing using time interval production tasks (e.g., Jazayeri & Shadlen, 2010).

We hope REPP plays a major role in improving the efficiency, scalability, and reach of SMS research. Finding new ways to allow online data collection has become particularly important during the COVID-19 pandemic, with many researchers unable to run experiments in the laboratory. Supporting online experiments on SMS will also significantly reduce the time and resources that researchers usually spend to recruit and test participants in the laboratory. Moreover, online experiments allow for the collection of significantly larger and more diverse samples of participants, both demographically and culturally. This is crucial for moving away from the relatively restricted and small samples of university students that laboratory studies tend to rely on (Henrich et al., 2010). Since online SMS experiments can be more accessible and easy to share, they can also increase research diversity and collaboration worldwide, an important challenge in today’s cognitive science (Barret, 2020). Finally, by enabling online SMS experiments, REPP opens new avenues for research on SMS that would be nearly impossible in the laboratory. For example, REPP has been previously used to collect large tapping datasets to study individual differences on SMS in the first GWAS study on beat synchronization (Niarchou et al., 2021). Similarly, the ability to collect large tapping datasets online can help increase our understanding of the role of SMS in the context of various neurodevelopmental disorders, including attention deficit hyperactivity disorder (Noreika et al., 2013), dyslexia (Colling et al., 2017; Thomson & Goswami, 2008), and Parkinson’s disease (Bieńkiewicz & Craig, 2015). Alternatively, experiments may recruit large groups of participants from disparate cultural backgrounds to better understand the cultural foundations of SMS and auditory perception (Jacoby et al., 2020). REPP makes this possible while massively increasing the reach, scalability, and speed of data collection.

## Acknowledgements

This technology is named after Dr. Bruno Repp, notable for his work on sensorimotor synchronization, rhythm, and timing perception. His pioneering research has been truly inspirational. We also thank the members of the Computational Auditory Perception group for their help and feedback, as well as the laboratory staff in Max Planck Institute for Empirical Aesthetics for their assistance during data collection.

## Appendix A Detailed methods

## Implementation

We implemented REPP as a *Python* package. In all experiments, REPP was integrated into our in-house system to perform behavioural experiments, PsyNet (Harrison et al. 2020). This system is based on the Dallinger framework^2^ for hosting and deploying experiments. Participants interact with the experiment via a web browser, which communicates with a back-end Python server cluster responsible for organizing the experiment and communicating with REPP. This cluster can run using a local webserver (for in-lab experiments) or by a cloud Platform as a Service such as Heroku (for online experiments). Currently, PsyNet is only supported by Google Chrome.

## Instructions

When using REPP, participants should be informed that the experiment can only be performed using laptop speakers (e.g., do not use headphones or wireless devices). We also suggest using a volume calibration test to adjust the volume of the speakers to a level that is sufficiently good to be detected by the microphone. In our experiments, we used a volume calibration test to play an audio stimulus through the speakers and record the signal with the built-in microphone, using a sound meter to visually indicate whether the level was appropriate or not (see Figure S1 for a screenshot of the volume test). Finally, participants should be clearly instructed about how to tap on their laptop in a way that is compatible with REPP and also feels natural to them: “Tap on the surface of your laptop with your index finger (e.g., do not tap on the keyboard or tracking pad, and do not tap using your nails or any object)”. Here we used a tapping calibration test to ask participants to practise tapping in the required way and test if the microphone could detect their signal, also using a sound meter to give feedback visually (see Figure S1 for a screenshot of the tapping test). In cases where the signal was too low, participants were indicated to tap in different locations of the laptop or try to tap louder.

## Pre-screening Tests

When running experiments online, it is important to ensure participants follow the instructions and perform the task as required (e.g., Clifford & Jerit, 2014; Crump et al., 2013). In addition, REPP has several technical requirements that participants must meet in order to provide valid tapping data. To address this, we used two pre-screening tests in the online experiments reported in this paper (Experiment 3 and 4): an attention test and a recording test.

## Attention Test

The attention test was used to determine whether participants were paying attention to the instructions or not (see Figure S2 for a screenshot of the attention test). The test consisted of two pages that could only be passed if a participant carefully read the instructions. The attention test was presented at the beginning of the experiment after asking for general demographic information. In our implementation, participants who failed the first page in the attention test were excluded from the experiment, whereas the second page was used for post-hoc quality assessment (we did not exclude participants based on failure to answer correctly in the second page).

## Recording Test

The recording test was used to determine whether participants were using hardware and software that did not meet the technical requirements of REPP, such as malfunctioning speakers or microphones, or the use of strong noise-cancellation technologies (see Figure S3 for a screenshot of the recording test). The recording test was used at the beginning of the experiment, after providing general instructions with the technical requirements of the experiment. Thus, this test can also be used as an attention test, as participants must follow the given instructions (e.g., accept the enabling of the microphone in the browser, unplug any headphones or wireless devices, turn up the volume of the computer) in order to successfully pass the test. For example, bots that click randomly on the screen would naturally not be able to complete these steps. The recording test consisted of a recording page that played a test stimulus with six marker sounds. The markers were recorded with the laptop’s microphone and analyzed using the signal processing pipeline. During the marker playback time, participants were supposed to remain silent (not respond). In our implementation, we used two recording trials. Those cases in which all marker sounds could not be detected in one of the two recording trials were excluded from the experiment.

**Figure S1.**
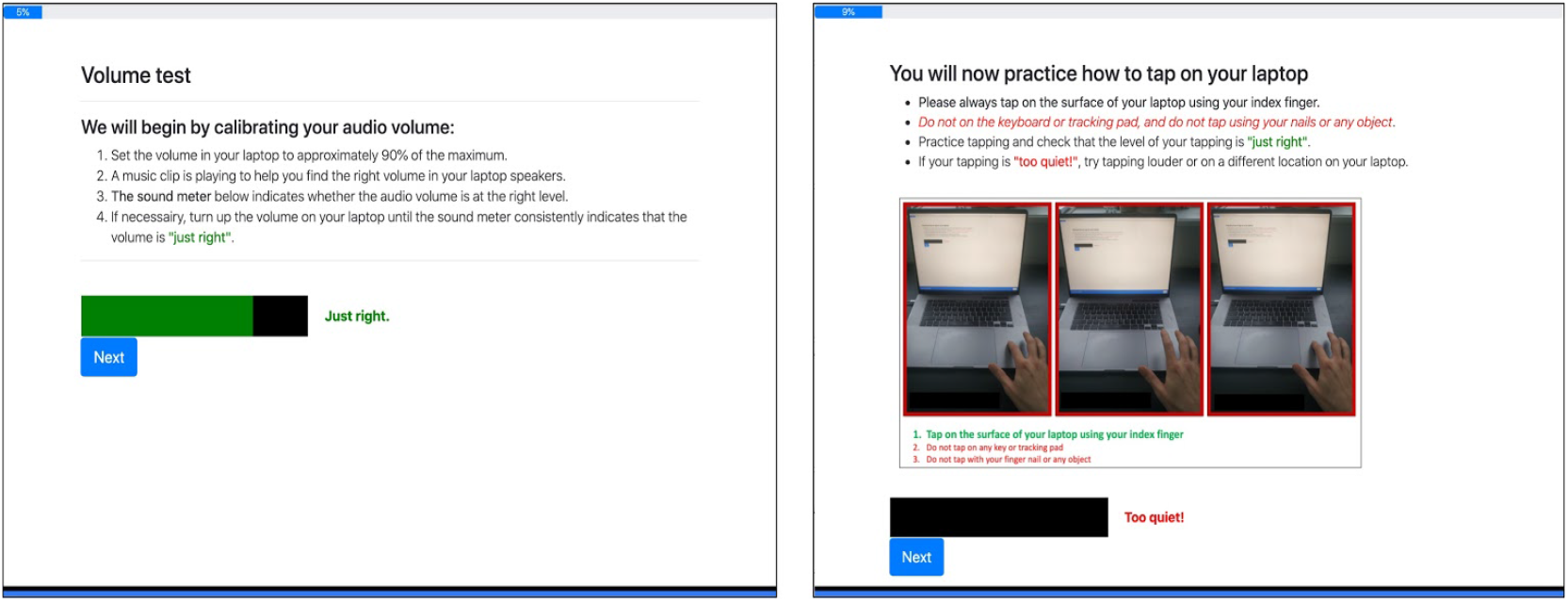
Volume and tapping calibration tests using sound meters to provide visual feedback.

**Figure S2.**
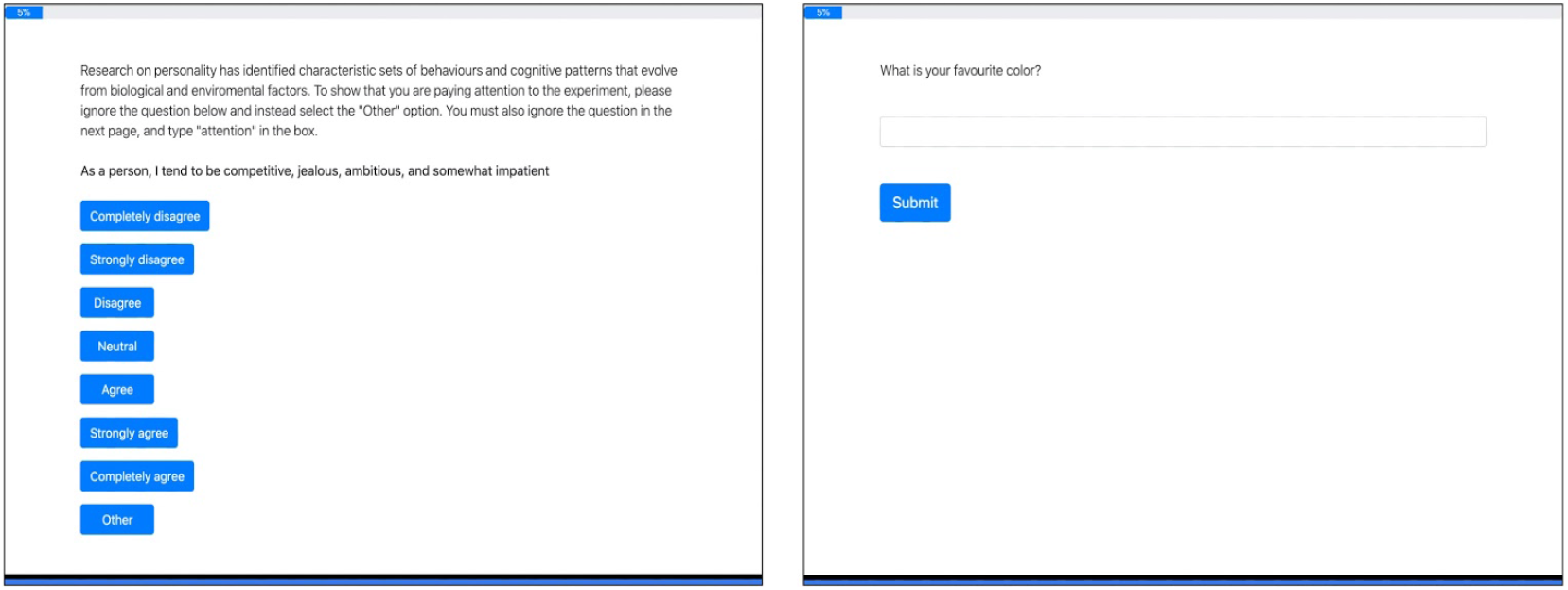
Attention test to determine whether participants follow the instruction in online experiments.

**Figure S3.**
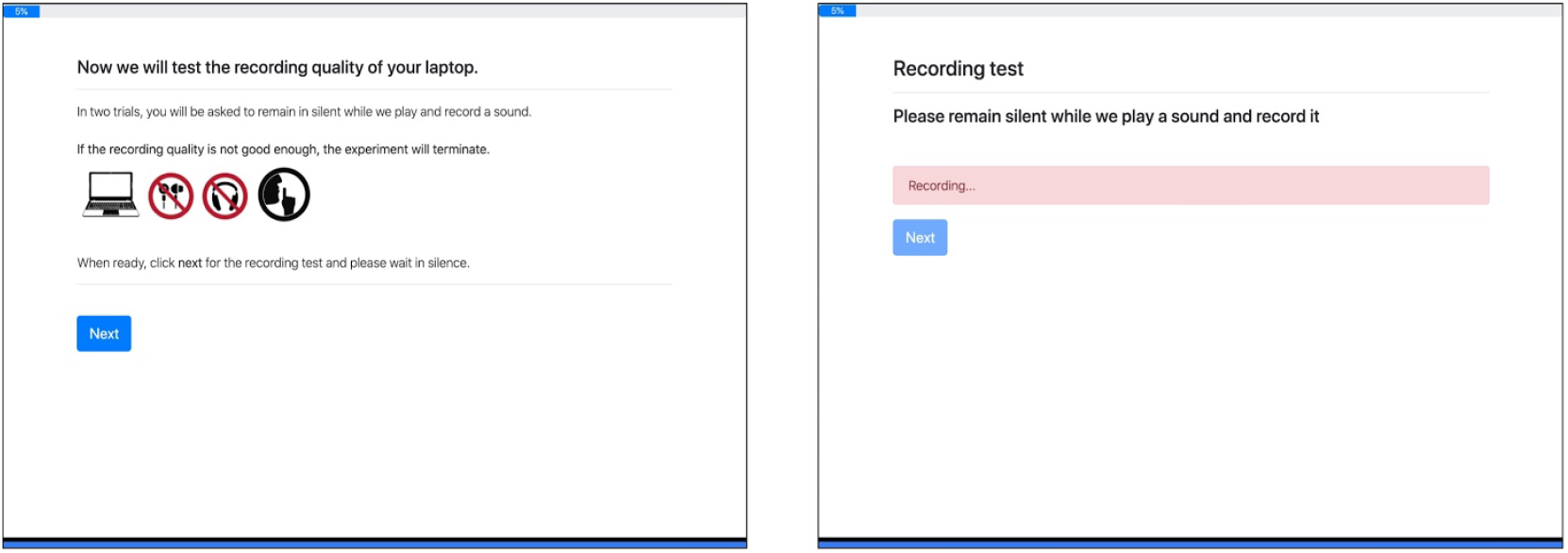
Recording test to determine the performance of REPP in online experiments.

## Practice Phase

In Experiment 2 (laboratory), the practice phase consisted of two trials of isochronous tapping to a metronome sound (each trial was 20 seconds long, one with IOIs of 800 ms and the other with IOIs of 600 ms). Participants performed a practice phase the first time they used each method, one for REPP and one for the independent in-lab method. A researcher was present during the practice phase to provide feedback on participants’ practice trials.

In experiment 3 (online), the practice phase consisted of four trials of isochronous tapping to a metronome sound (two with IOIs of 800 ms and two with IOIs of 600 ms, 20 seconds long each). In experiment 4 (online), the practice phase consisted of four trials using the stimulus sampling procedure used in the main experiment (e.g., three-interval rhythms randomly sampled from the triangular simplex with a fixed duration of 2,000 ms and repeated 10 times). In the two online experiments, the recording of the first practice trial was analyzed in real time to provide feedback based on the quality of the audio and tapping signal (using the *Failing Criteria* described below). If the signal of the recording did not pass the failing criteria, participants were reminded of the instructions and were able to continue with the other practice trials. At the end of the practice phase, all trials were analysed online using the same procedure and participants who failed two or more trials were excluded from the experiment. All participants were compensated proportionally to the time spent in the experiment, even if they failed the screening tests or practice phase. Figure S4 shows an example of the instructions and tapping trial given in the practice phase of Experiment 3. In each trial, to help participants only tap when the stimulus was played (and remain silent when the marker sounds were presented), we visually indicated on the screen when to start and when to stop tapping.

**Figure S4.**
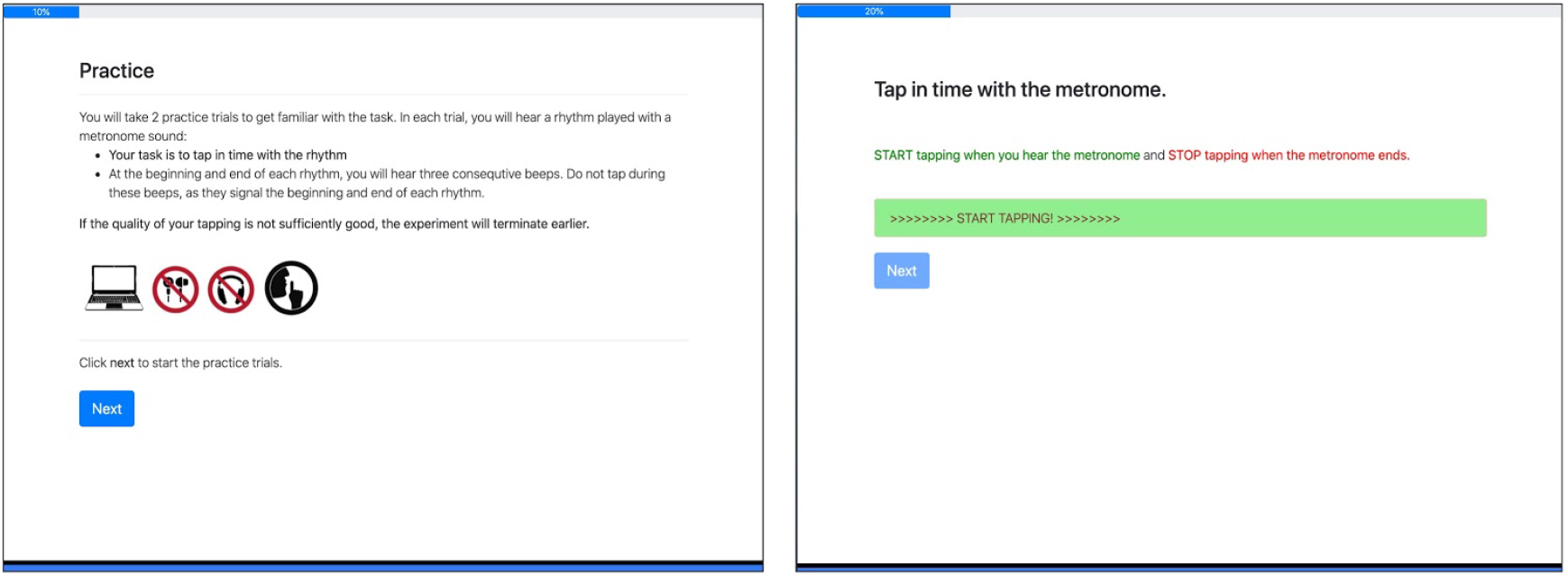
Instructions and tapping trial in the practice phase.

## Beat Synchronization Task

In the beat synchronization task (Experiment 2 and 3), participants were instructed to tap in time to with the beat until the music ends (Figure S5 shows a screenshot of the instructions for the beat synchronization task). As commonly used in this type of paradigm, to help participants find the beat, a metronome marking the beats in the first 11 seconds of the clip was added to the stimulus. To motivate participants to continue tapping accurately until the end of the clip we also added three more metronome beats to the end of the recording. Thus, to calculate participants’ tapping performance in this task, we only analysed the stimulus onsets when the metronome was not played. The materials of the beat synchronization task consisted of four 30-second long excerpts of music from two distinct music genres with different style, tempo, and tapping difficulty: track 1 (“You’re the First, the Last, My Everything” by Barry White) and track 2 (“Le Sacre du Printemps” by Stravinsky). The presentation order was fixed, namely: track 1, track 2, track 1, and track 2. The musical excerpts were taken from the MIREX 2006 Audio Beat Tracking database, which also provides annotations for beat locations given by listeners who tapped along to the music (McKinney et al., 2017). Based on these annotations, we identified the target beat locations from those consistently produced by the annotators using the following procedure: First, we performed kernel density estimation with a kernel width of 20 ms to find the mode of participants’ responses in any given time. Second, we locate the peaks of the probability density to find all onset locations in the music by identifying local maxima in the kernel density function.

**Figure S5.**
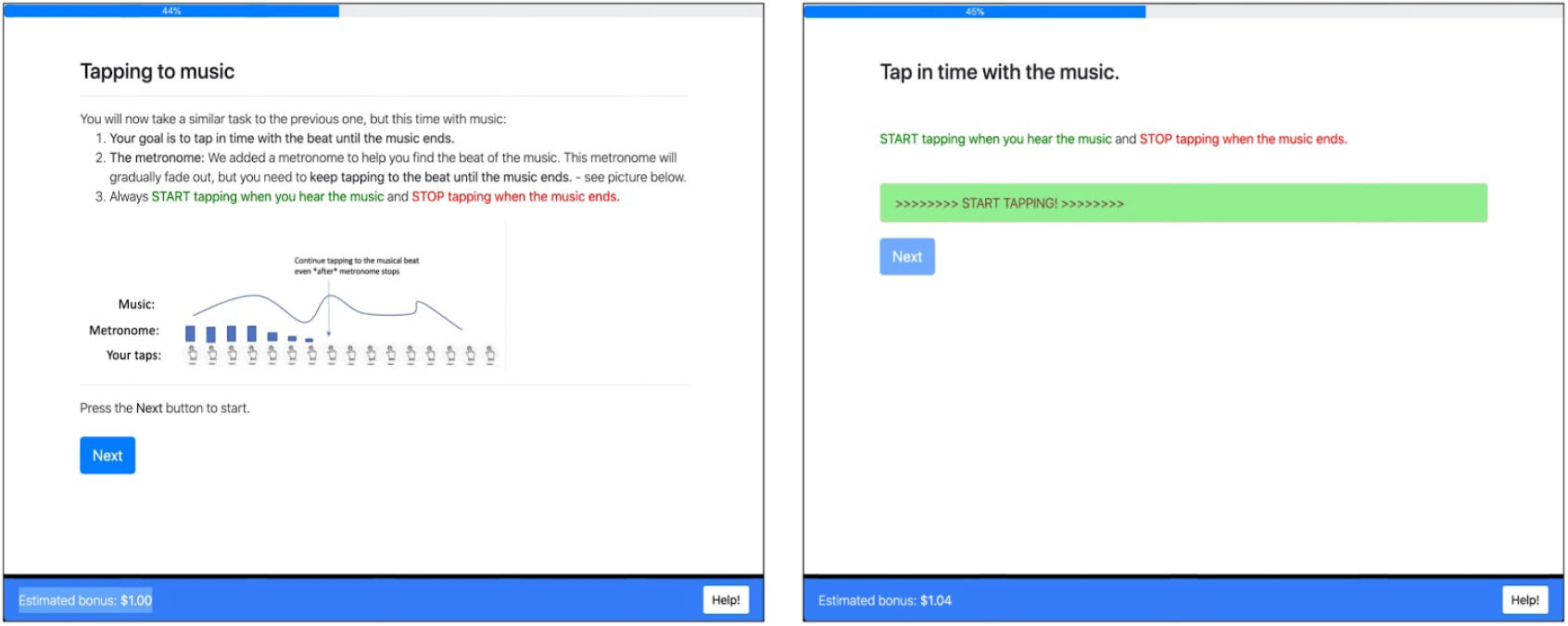
Instructions for the beat synchronization task.

## Failing Criteria

When measuring SMS in online experiments, it is crucial to determine whether participants are tapping in the required way (e.g., following the instructions) and whether any technical constraints may preclude the recording of their signal, such as cases with poor internet connection, malfunctioning hardware, or strong noise-cancellation technologies. To identify and exclude these cases in the online experiments reported in this paper (Experiment 3 and 4), we used two-step failing criteria. First, since REPP cannot work efficiently unless it detects all marker sounds with high precision, we failed all trials in which we could not detect all marker sounds included in the stimulus preparation step, or where the markers were displaced relative to each other for more than 15 ms. Second, we failed all trials where the percentage of detected taps (i.e., the number of detected tapping onsets out of the total number of stimulus onsets) was less than 50% or more than 200%. This measure is useful to deter participants from not responding at all or from tapping at an extremely fast rate, irrespective of the audio stimuli. Importantly, none of these criteria exclude trials based on actual tapping performance, but only based on whether the signal can be correctly recorded and processed by REPP and whether participants produced a minimally/ maximally acceptable number of tapping responses.

In experiment 3 and 4, the failing criteria was used in the practice phase to exclude participants who did not provide at least two valid tapping trials. We also used the failing criteria in the main tapping tasks to fail individual tapping trials. Moreover, as a data cleaning step, we removed from the analysis all tapping trials where the markers were displaced relative to each other for more than 5 ms, ensuring that we only included cases with nearly optimal latency and jitter.

## Participants

All participants provided consent in accordance with the Max Planck Society Ethics Council approved protocol (application 2018-38).

## Experiment 2

Participants were recruited using the internal database of the Max Planck Institute for Empirical Aesthetics (Frankfurt, Germany), with the requirement that they were at least 18 years old and had a basic understanding of English. The experiment took about 1 hour and the reimbursement was 14 €. A total of 20 participants (10 female, 10 male), aged 20-59 (*M* = 30.05, *SD* = 11.88) took part in the experiment.

## Experiment 3

All participants were recruited online using Amazon Mechanical Turk. We asked for five requirements in order to take part in the experiment: (i) participants must be at least 18 years old, (ii) participants must be fluent English speakers, (iii) participants must use a laptop to complete the experiment (no desktop computers allowed), (iv) participants must use an up-to-date Google Chrome browser (due to compatibility with PsyNet), and (iv) participants must be sitting in a quiet environment (to ensure that their tapping could be recorded precisely). In addition, to help recruit reliable participants, we only recruited participants with a 95% or higher approval rate on previous tasks on Amazon Mechanical Turk. Participants were paid at a US $9/hour rate according to how much of the experiment they completed (e.g., if participants failed a pre-screening task and left the experiment early, they were still paid proportionally for their time). The complete experiment took approximately 20-25 minutes.

A total of 226 participants provided valid tapping data in at least one trial, having already excluded all those who failed the pre-screening tests or the practice phase. For those participants who reported demographic information, ages ranged from 19 to 77 (*M* = 35.9, *SD* = 11.9), and 46% identified as female (54% as male).

## Experiment 4

Participants were recruited online using Amazon Mechanical Turk and the same requirements described in Experiment 3. The complete experiment took approximately 15-20 minutes. A total of 157 participants provided valid tapping data in at least one trial, having excluded those who failed the pre-screening tests and practice phase. For those participants who reported demographic information, ages ranged from 18 to 69 (*M* = 36.4, *SD* = 9.97), and 51% identified as female (48% as male and 1% as other).

## Appendix B Trialling mean asynchrony as an alternative measure to SD of asynchrony

Here we repeat the main analyses concerning tapping accuracy (Experiment 2 and 3) using mean asynchrony instead of *SD* of asynchrony (see Figure S6). Overall, the results are very similar to the ones reported in the text using *SD* of asynchrony (see Figure 3 and 4, respectively).

**Figure S6.**
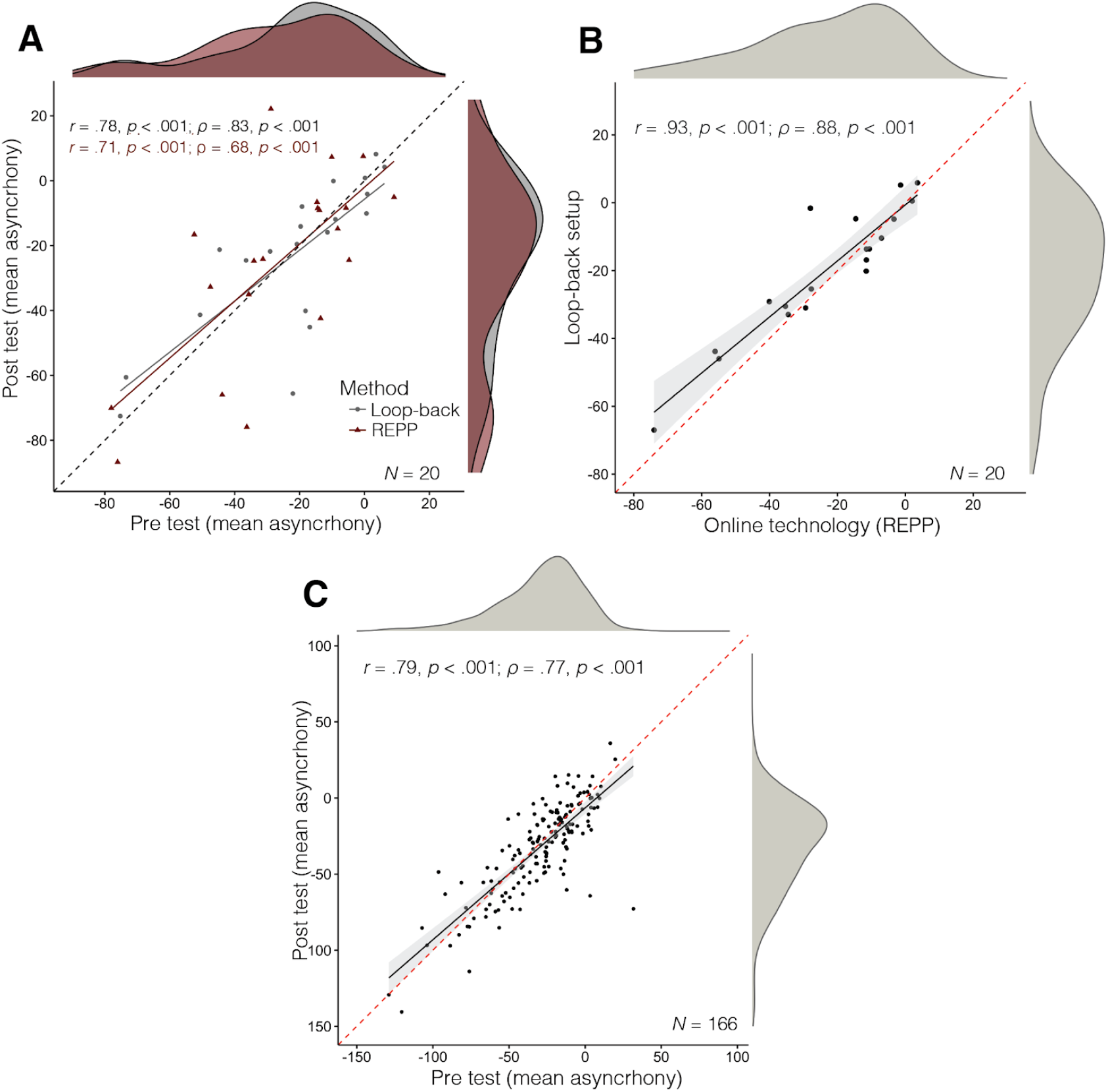
Replication of tapping accuracy analysis using mean asynchrony. (A) Experiment 2: test-retest reliability measured in the two methods. (B) Experiment 2: concurrent validity of REPP. (C) Experiment 3: test-retest reliability of REPP when measuring participants’ tapping performance online.

## Footnotes

1 There are two main methodological differences between the original experiment and the online replication: (i) the original experiment consisted of longer experimental sessions with few participants (*N* = 14; with 20 to 30 full chains per participant), and (ii) it did not mix the order of the iterations from different chains (participants had to complete a single chain before moving to the next one). In contrast, the online replication consisted of shorter sessions with more participants and mixed the order of the iterations from different chains.

2 https://github.com/Dallinger/Dallinger

